# Tactile pup loss and acoustic signal enhance selective maternal retrieval behavior in echolocating bats, *Pipistrellus abramus*

**DOI:** 10.64898/2026.01.08.698298

**Authors:** Shizuki Nara, Kazuki Yoshino-Hashizawa, Kohta I Kobayasi, Shizuko Hiryu

## Abstract

Pup retrieval is a critical maternal behavior ensuring pup survival. Its mechanisms have been studied mainly in rodents, where mothers tolerate brief pup separation. In contrast, in species for which even short separation is life-threatening, mothers maintain constant physical contact, reinforcing anti-separation behavior. Studying such species may uncover maternal strategies under survival-critical extremes. Among these species, echolocating bats provide an ideal model for acoustic-based decision-making, as they rely heavily on sound. In bats, pup isolation calls (ICs) are essential signals in mother-pup communication. To assess their motivation in retrieval based on ICs, we conducted behavioral experiments: (1) pup retrieval test, and (2) playback test using ICs from own vs. non-own pups. Each was conducted under three maternal conditions: holding (a) no-pup, (b) one-pup, and (c) two-pup, to assess tactile influence on maternal motivation. In pup retrieval tests, mothers showed selective responsiveness to their own pups, and it is decreased with their pup development. Playback tests showed that acoustic signals alone were sufficient to elicit maternal responses, and their latencies were negatively correlated to pup vocalization rates. These maternal responses were strongly suppressed under condition (c), but robustly expressed under conditions (a) and (b). These findings suggest that maternal motivation is context-dependent responsiveness, which is enhanced by tactile pup loss and ICs, providing insight into instinctive maternal behavior in mammals.

**Summary statement:** This study demonstrates that in the *Pipistrellus abramus*, maternal motivation is context-dependent responsiveness, which is enhanced by tactile pup loss and vocalizations, providing insight into a flexible multimodal mechanism underlying instinctive maternal behavior.

## Introduction

Maternal behavior is essential for the survival of mammalian offspring, which are typically born in a highly altricial state. Parental care itself is widely observed across birds, reptiles, amphibians, fishes, and even invertebrates (Balshine and Sloman, 2011; Cockburn, 2006; Rosenblatt and Snowdon, 1996; Schulte et al., 2020) In mammals, species lacking maternal care—particularly nursing—do not exist (Kuroda et al., 2024). Maternal behavior in mammals represents one of the most stable and evolutionarily conserved species-specific motivated behaviors and plays a crucial role in reproductive success (Rosenblatt, 1967; Wang et al., 2022). A representative and essential component of maternal behavior is pup retrieval (Lonstein and Fleming, 2001). In rodents such as mice and rats, pup retrieval is defined as the act of returning displaced pups to the nest and has long been used as a primary index of maternal behavior research (Beach and Jaynes, 1956; Noirot, 1972; Numan and Insel, 2003). In these species, mothers periodically leave the nest to forage, resulting in brief periods during which mothers and pups are separated.

In contrast, in species in which mothers always carry their offspring during the early postnatal period, even brief periods of physical separation can be directly life-threatening for the pup. Therefore, in species of this kind, when mother-infant separation occurs for any reason, behaviors that enable the mother to rapidly retrieve her pup may have been selectively reinforced. The Japanese house bat, *Pipistrellus abramus*, which is the focus of the present study, exhibits precisely this life-history pattern. For several days after birth, mothers continuously carry their pups, and any separation during this early period poses a serious risk to pup survival. These observations suggest that the importance of retrieval as a component of maternal behavior is particularly high in *P. abramus*. It should be note that because retrieval in this species does not involve returning the pup to a nest, it is necessary to define pup retrieval in a manner different from the conventional definition used for rodents. Whereas pup retrieval in mice—according to the Stanford Mouse Ethogram—is defined as “the dam picking up a pup with her mouth and transporting it from outside to inside the nest,” pup retrieval in echolocation bat, *Rhinolophus ferrumequinum* was defined following Matsumura (1979) as “the mother approaching and taking up a pup that has become separated from her,” referring specifically to the moment the pup is reattached to the mother.

*Pipistrellus abramus* is an echolocating bat species and a mammal that relies heavily on auditory information (Hiryu and Riquimaroux, 2011; Yoshino-Hashizawa et al., 2023). This strong dependence on acoustic cues makes it an excellent model for evaluating decision-making based on quantifiable vocal signals. In general, newborn bats produce both precursor forms of echolocation calls and vocalizations known as isolation calls (Hiryu and Riquimaroux, 2011; Jones et al., 1991). Isolation calls, in particular, are produced only during the early postnatal period and are thought to function as signals soliciting maternal care (Engler et al., 2017; Gould, 1975). Many bat species form maternity colonies consisting of dozens of individuals and rear their young in groups (Funakoshi et al., 2009; McCracken and Wilkinson, 2000). Given their strong reliance on auditory information, isolation calls are believed to provide crucial cues that enable mothers to recognize their own offspring. Indeed, in species such as the Mexican free-tailed bat (*Tadarida brasiliensis mexicana*), isolation calls have been shown to contain acoustic features that allow individual identification (Knörnschild and Von Helversen, 2008).

In bats, singleton births are the most common reproductive strategy (Garbino et al., 2021). Multiple births, in which a female produces more than one pup per pregnancy, have been reported only in a limited number of species, primarily within the family Vespertilionidae (Barclay et al., 2003). *Pipistrellus abramus* is one such species, producing 1–3 pups per year, with twins being the most common (Uchida, 1950). For *P. abramus* mothers, who constantly carry their pups, whether or not they are currently holding a pup is expected to influence their motivation to perform pup retrieval behavior. In this study, we aimed to clarify the conditions under which pup retrieval—one of the key components of maternal behavior—is expressed in *P. abramus*, and to identify the factors that elicit this behavior, particularly how pup isolation calls and the physical state of mother-infant contact affect maternal responsiveness. To this end, we conducted behavioral experiments in an indoor experimental arena, presenting mothers with pups aged 3–13 days or with pup vocalizations, and quantitatively measured the extent to which retrieval behavior was expressed.

## Materials and Methods

### Study species and ethical status

The study species was the Japanese house bat (*Pipistrellus abramus*), a member of the family Vespertilionidae. Females of this species give birth once a year, usually to two or three pups, and in the wild form colonies of 20–30 individuals that communally raise offspring. Pregnant females were captured from a colony roosting near the Doshisha University campus in Kyotanabe, Japan, and subsequently housed in the laboratory. Each female was placed in a plastic cage (370 × 220 × 240 mm) equipped with a paper dome roost (146 × 98 × 54 mm) and provided ad libitum access to mealworms and water. Mothers gave birth between late June and early July, and pups were reared together with their mothers in the same cage.

In total, 16 mothers and 37 pups were used as subjects in this study. The pups comprised 19 males and 18 females. The experiments were conducted over three breeding seasons from 2023 to 2025, involving 6 mothers and 12 pups in 2023, 5 mothers and 13 pups in 2024, and 5 mothers and 12 pups in 2025. Eight mothers gave birth to three pups, seven mothers to two pups, and one mother to a single pup. Two pups died shortly after birth, and the remaining 37 pups were used in the study.

All experiments complied with the Principles of Animal Care (NIH publication no. 86-23, revised 1985) and the relevant Japanese laws, and were approved by the Animal Experiment Committee at Doshisha University (Permission No.: A25021, A24018, A23018).

### Experiment environment

All experiments were conducted in a sound-attenuated chamber (1.4 × 1.4 × 2.1 m) illuminated with red light (G-82H(R); ELPA, Japan) to minimize the use of visual cues. At the center of the chamber, a cylindrical arena (70 cm in diameter, 50 cm in height) enclosed by a metal mesh wall was installed. Within the arena, a plastic-walled observation box was placed for behavioral tests (20 × 10 × 10 cm for Experiment I and 30 × 20 × 10 cm for Experiment II), where mother-pup interactions were recorded.

The behavior of mothers and pups was monitored using an infrared camera (DMK 33UX2733; The Imaging Source, Germany; frame rate: 10 fps) positioned 70 cm above the arena. Vocalizations were recorded with an ultrasonic microphone (Anabat SD2 CF Bat Detector; Titley Scientific, Australia) placed on the arena floor outside the observation box (Figs. 1 and 2). Acoustic signals were digitized using a data acquisition system (BNC-2110 / PXIe-6358 / PXIe-1073; NI, USA) controlled by a custom LabVIEW 2016 program (NI, USA; sampling rate: 500 kHz/channel). The recorded vocalizations were later analyzed to quantify developmental changes in the number of isolation calls produced by pups across postnatal age. Pup skin surface temperature was measured through the fur immediately after separation from the mother while the pup was freely moving. Measurements were obtained using a thermographic camera (Xi400; Optris, Germany) positioned 70 cm above the arena and controlled by dedicated software (PIX-Connect; Optris, Germany).

**Fig. 1.**
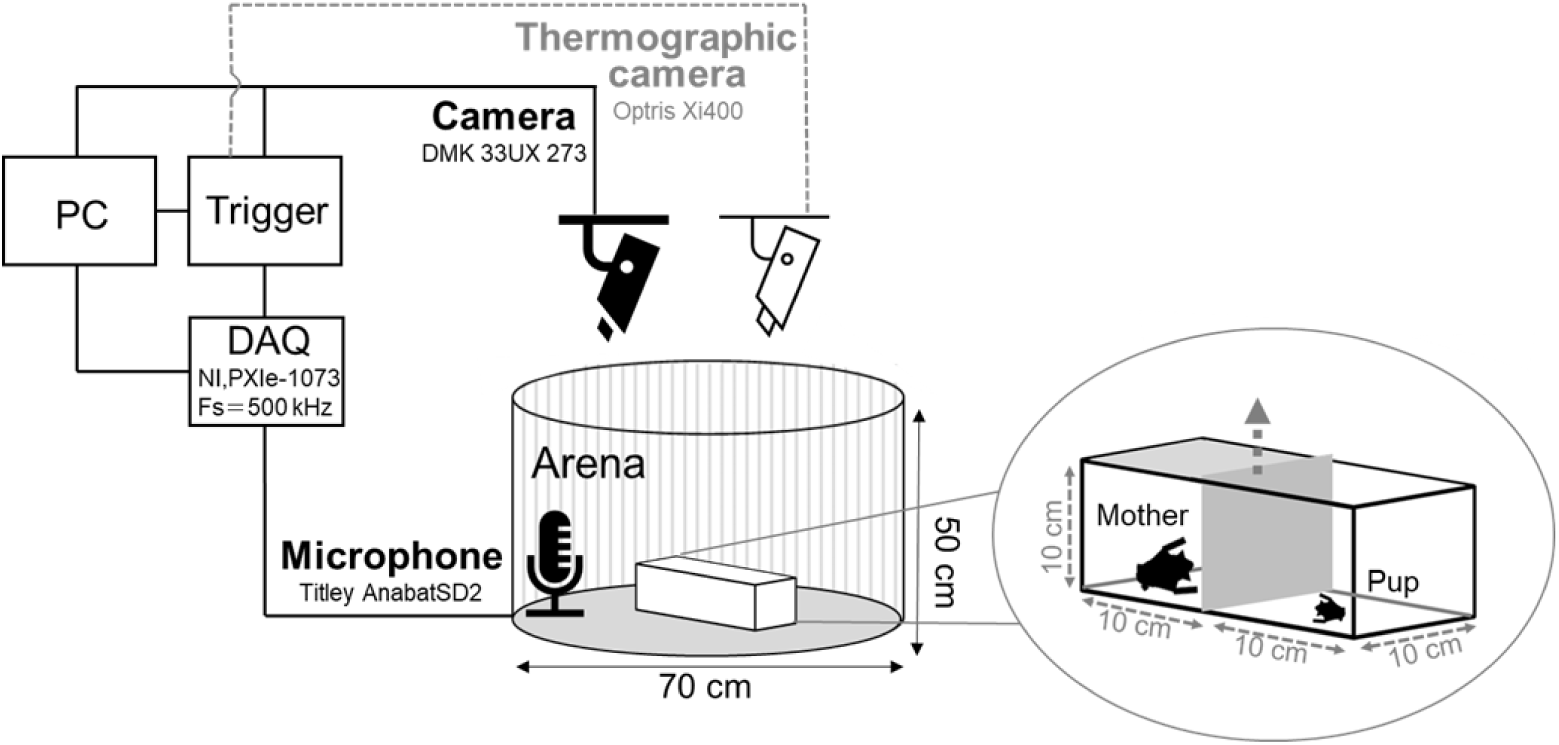
Experimental setup for the pup retrieval test (Experiment I). The test arena consists of two compartments separated by a removable partition. The mother’s compartment was enclosed by opaque walls to block visual and echolocation cues, whereas the pup’s compartment remained open at the top. A retrieval event was defined as the mother approaching and picking up her pup after the partition was removed.

**Fig. 2.**
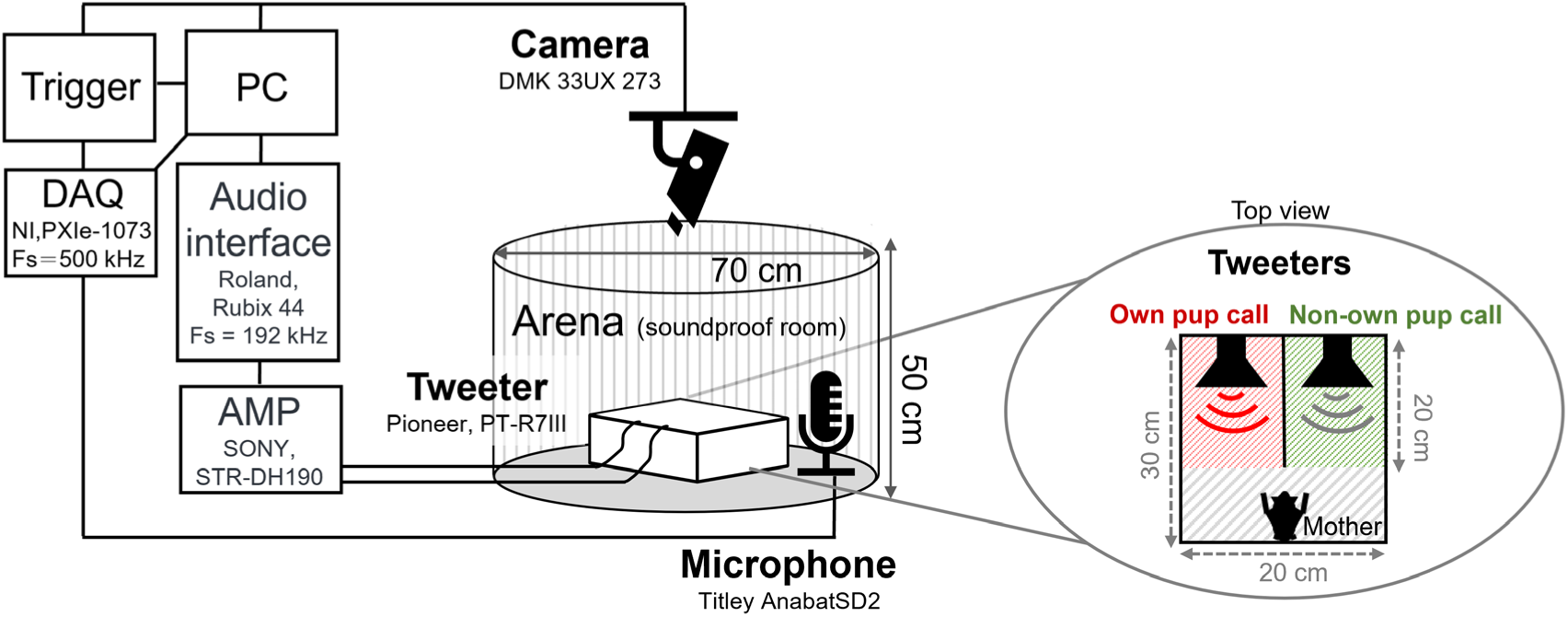
Experimental setup for the playback test (Experiment II). The observation box was divided into three zones: two playback zones equipped with tweeters broadcasting calls from the mother’s own and a non-own pup, and a central waiting zone where the mother was initially placed. Maternal choice behavior and latency to initiate a response (only under the no-pup condition) were recorded.

In this study, we defined the maternal behavior of approaching and retrieving a pup as retrieval (Matsumura, 1979), and conducted two experiments to test whether pup-derived stimuli elicited this response.

### Experiment I: Pup retrieval test

Maternal retrieval behavior was observed in 11 adult females. The Pup Retrieval Test (PRT) was conducted every other day when their pups were 3-13 days old, and additionally on postnatal days 20 and 30 (under the no-pup condition). The observation box was divided into two compartments (10 × 10 × 10 cm each): one for the mother and the other for the pup (Fig. 1). The compartments were separated by a removable partition lined with sound-absorbing material. The mother’s compartment was enclosed by opaque plastic walls, including the top, to prevent visual and echolocation-based access to the surrounding area, whereas the pup’s compartment remained open at the top.

Each mother, holding all of her pups, was allowed to habituate to the observation box for 10 min. After habituation, the experimenter gently detached each pup from the mother and placed in the pup compartment, while the mother remained in her compartment. The experimental conditions and the number of individuals tested in each condition are summarized in Table S1. Infrared cameras, microphones and thermographic camera were synchronized by a trigger signal, and recording began simultaneously with the removal of the partition between the two compartments. Each trial ended when the mother reached her pup or 15 min had elapsed. After each trial, the mother and her pup were reunited and returned to their home cage.

Each mother was first tested with her own pup(s). Mothers that gave birth to twins were tested twice (once with each pup), and those that gave birth to triplets were tested three times, once for each pup, on the same experimental day. For four mothers that consistently retrieved their own pups, additional trials were conducted using pups born to other females to compare retrieval responses toward own versus non-own pups (Table S2, total 9 trials).

To investigate whether physical contact with pups affected retrieval motivation, mothers were tested under three conditions (Table S1): (a) no-pup condition, in which mothers were tested without holding any pups (n = 6 mothers, total 84 trials; all conducted in 2023, with all pups removed), (b) one-pup condition, in which mothers held one pup during the test (n = 2 mothers, total 24 trials; all trials in 2024, mothers that had given birth to twins, with one pup temporarily removed), and (c) two-pup condition, which mothers held two pups during the test (n = 3 mothers, total 54 trials; all trials in 2024, mothers that had given birth to triplets, with one pup temporarily removed). During all trials, the non-holding littermates were kept with their littermates in a separate enclosure.

To examine factors affecting pup retrieval behavior, we employed generalized linear mixed models (GLMMs) with a binomial error distribution and a logit link function. The response variable was whether the mother retrieved the pup (retrieved vs. not retrieved). The number of pup holdings and the postnatal age of the presented pup were included as fixed effects. Mother identity was included as random effect to account for repeated observations from the same individual. Model fitting was conducted using the glmer function in the lme4 package in R (ver. 4.5.0). To assess the efficacy of the explanatory variables, candidate models were compared using Akaike’s Information Criterion (AIC). The statistical significance of the fixed effects was assessed using Wald z-tests based on the model with the lowest AIC.

### Experiment II: Playback test

In Experiment I, odors and other sensory cues were not excluded. Therefore, to examine whether mothers used pup vocalizations as cues for retrieval behavior, we conducted a playback test (Experiment II), in which the available cues were restricted to acoustic information only. In Experiment II, we controlled for potential effects of odor and pup movement that were not accounted for in Experiment I. Five mothers were tested: one that had a single pup, two that had twins, and two that had triplets. The playback test was conducted on alternated days when pups were 3–13 days old, as in Experiment I. The experimental setup was similar to that used in Experiment I. An observation box (30 × 20 × 10 cm) was placed inside the arena and divided into three zones by plastic partitions (Fig. 2). The two side zones each contained a tweeter (PT-R7Ⅲ; Pioneer, Japan), and the mother was placed in the central waiting zone (20 × 10 × 10 cm). The mother’s own pup’s call, recorded on the same day as the experiment, was broadcast from one tweeter, while the call of a non-own pup (matched in postnatal age but mismatched in sex) was simultaneously broadcast from the other. The left–right tweeter assignment was randomized across trials to prevent side bias, and audio-video recording began at the same time. The trial ended when the mother entered either playback zone (choice) or when 5 min had elapsed. The mother’s choice of zone was recorded to determine whether she preferentially responded to her own pup’s calls. Figure 3 shows representative examples of pup calls used for playback. Each mother was presented with calls from each of her own pups across experimental days in a balanced order within litters. After the trial, the mother was returned to her home cage with her pups.

**Fig. 3.**
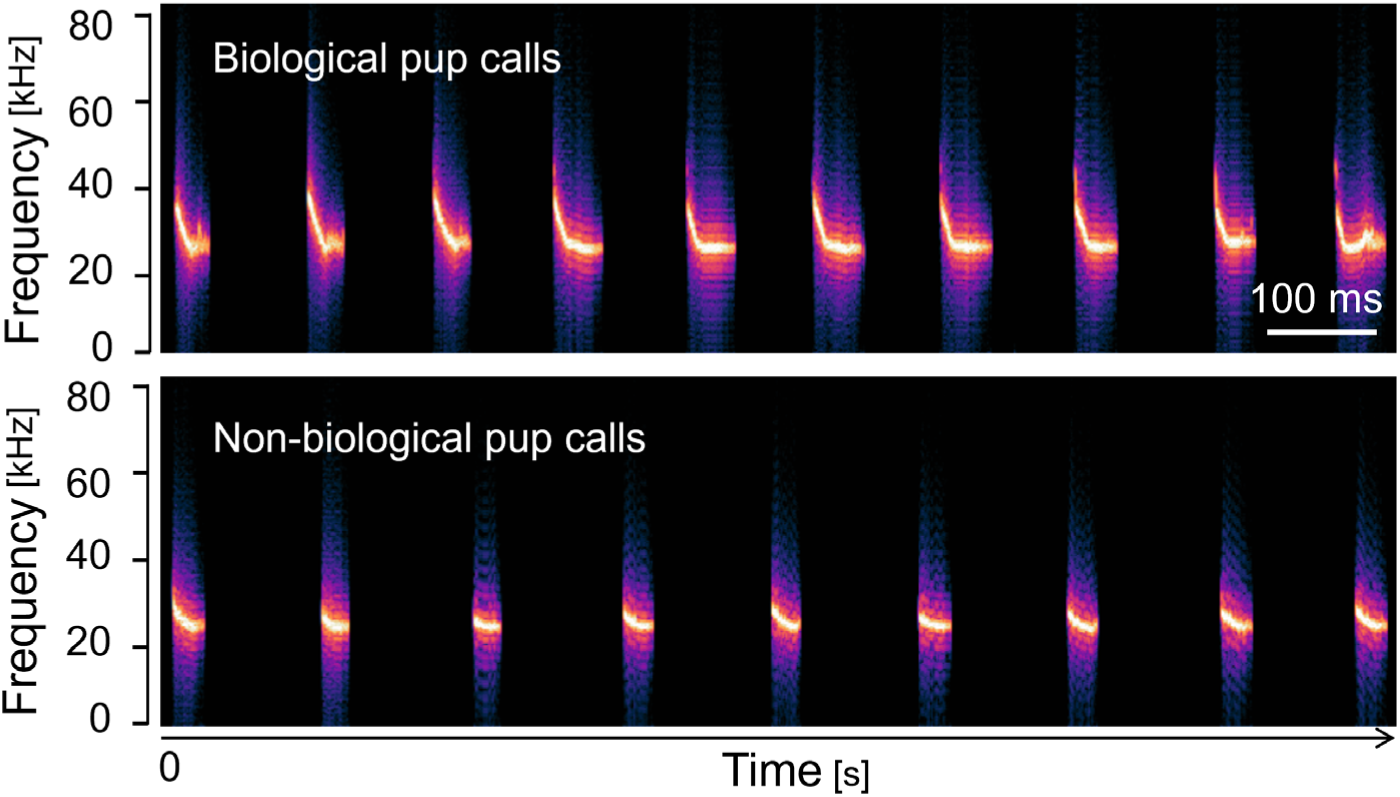
Representative spectrograms of pup isolation calls used in the playback experiments. Spectrograms show the frequency modulation pattern and duration typical of day-7 isolation calls. The vertical line at 0 s indicates the onset of playback stimulation.

On each test day, pup vocalizations to be used for playback were recorded in advance. A pup was gently removed from its mother and placed in an open-top plastic box (20 × 20 × 20 cm), and the vocalizations were recorded using an ultrasonic microphone (Anabat SD2 CF Bat Detector; Titley Scientific, Australia) positioned approximately 10 cm above the pup. Playback stimuli were processed using a MATLAB-based denoising algorithm (Okabe et al., 2024) to remove environmental noise. For calls containing multiple harmonics, only the fundamental frequency component was retained for playback. To standardized playback amplitude, the processed sound was replayed through a tweeter and recorded by a second microphone positioned 10 cm away. The output voltage of the playback system was then adjusted to match the amplitude of the pup’s original calls. Each playback sequence lasted 30 s and was repeated continuously throughout the trial. The processed sounds were played through the tweeters using Audacity software (version 3.3.3, Muse Group) on a PC connected to an audio interface (Roland Rubix 44; sampling rate 192 kHz) and a stereo amplifier (Sony STR-DH190).

To determine whether the mothers responded merely to isolation calls in general, or specifically to calls that matched the pups’ current developmental stage, we conducted an additional playback experiment. Playback stimuli consisted of vocalizations, including isolation calls, recorded from pups at postnatal day 7, and the tests were performed on mothers whose own pups no longer produced isolation calls (postnatal days 20 (range: 17–20) and 35 (range: 32–35: owing to growth-dependent variability in experimental schedules)). Pup vocalizations containing isolation calls from her own pup and from a non-own pup (both recorded at day 7) were played simultaneously from the two tweeters, and video recording began at the same time. This additional playback experiment was conducted using the same procedures as the playback experiment described above.

As in Experiment Ⅱ, mothers were also tested under the three conditions (Table S3): (a) no-pup condition; mothers tested without holding any pups (n = 5 mothers, total 26 trials; all tested in 2025, with all pups removed), (b) one-pup condition; mothers holding one pup during the test (n = 2 mothers, total 11 trials; mothers that had given birth to twins, with one pup temporarily removed), and (c) two-pup condition; mothers holding two pups during test (n = 3 mothers, total 10 trials; mothers that had given birth to triplets, with one pup temporarily removed). Sample sizes varied across days and conditions, reflecting mortality and day-to-day variation in the availability of individuals suitable for testing.

A response score was assigned based on the mother’s behavior: entering a playback zone was scored as 1, whereas remaining in the waiting zone throughout the trial was scored as 0. To further assess behavioral selectivity, maternal responses were categorized according to whether the mother approached the zone broadcasting her own pup’s call, the zone broadcasting non-own pup call, or remained in the waiting zone (stay). The latency from playback onset to the mother’s first entry into a playback zone was also measured from the video recordings, but only under the no-pup condition (n = 4 mothers, total 20 trials; M-13 to M-16 in Table S3). When mother did not move between zones, the latency would be set to 300 seconds, the maximum measurement time. In echolocating bats, the number of emitted echolocation calls provides useful information for inferring decision-making processes. Therefore, we also quantified the mother’s echolocation call rate from the recorded audio.

From the playback sounds, pup vocalizations with a minimum frequency below 30 kHz were analyzed using USVSEG (Tachibana et al., 2020) to determine their duration. Following the developmental changes in pup vocal features previously reported for the Japanese house bat (*Pipistrellus abramus*), calls were categorized into long and short groups based on the duration distribution. The long group was defined as isolation calls (Hiryu and Riquimaroux, 2011), whereas the short group was classified as echolocation precursor calls. After classification, the number of isolation calls per second was calculated for each pup. In addition, the relative proportions of isolation calls and echolocation precursor calls were computed to quantify developmental transitions in vocal output.

To analyze maternal approach behavior, we employed GLMMs with a binomial error distribution and a logit link function. To analyze the overall approach behavior, the response variable was defined as the presence or absence of maternal approach toward the sound source (approach vs. no approach). The number of pup holdings and the postnatal day of the mother were included as fixed effects. Mother identity was incorporated as a random intercept, and in selected models, a random slope for the postnatal day of the mother was also included to account for individual variations in behavioral change across postnatal days.

To examine factors influencing the selection of pups after an approach, we analyzed whether mothers approached their own pups. In this analysis, we examined the impact of pup vocalizations by incorporating the call rates of both the subject pup and other pups in the environment or their differential as fixed effects. Mother identity was incorporated as a random intercept. We employed the glmer function of the lme4 package in R (ver. 4.5.0). The model selection was performed using Akaike’s Information Criterion (AIC). The statistical significance of the fixed effects was assessed using Wald z-tests.

## Results

### Experiment I: Pup retrieval test

The mothers exhibited retrieval behavior, approaching and picking up their own pups (see Supplemental Video 1). Figure 4 shows the proportion of trials in which retrieval was observed in 11 mothers, plotted against the pups’ postnatal age. In all three conditions, the retrieval rate was highest on postnatal day 3 (the first test day) and gradually decreased with age. Individual results for each mother are summarized in Table 1. Under the one-pup condition, retrieval behavior was no longer observed after postnatal day 9, and under the two-pup condition, it ceased after day 5. Even in the no-pup condition, retrieval behavior was absent by postnatal day 20, and the experiment was terminated on day 30 because the pups had begun to move independently, either leaving the arena or approaching the mother. Furthermore, when non-own pups were presented, mothers did not exhibit retrieval behavior, even under the no-pup condition (Table S2). Model comparison using GLMMs showed that additive model (without interaction between two fixed effects; the number of holdings and the pup age) had the lowest AIC (see Table S4 for all AICs). Both factors had significant negative effect on retrieval probability (Holdings: Estimate ± SE = -3.29 ± 1.47, *z* = -2.23, *p* = 0.026; Pup age: Estimate ± SE = -0.62 ± 0.26, *z* = -2.37, *p* = 0.018) in the selected model.

**Fig. 4.**
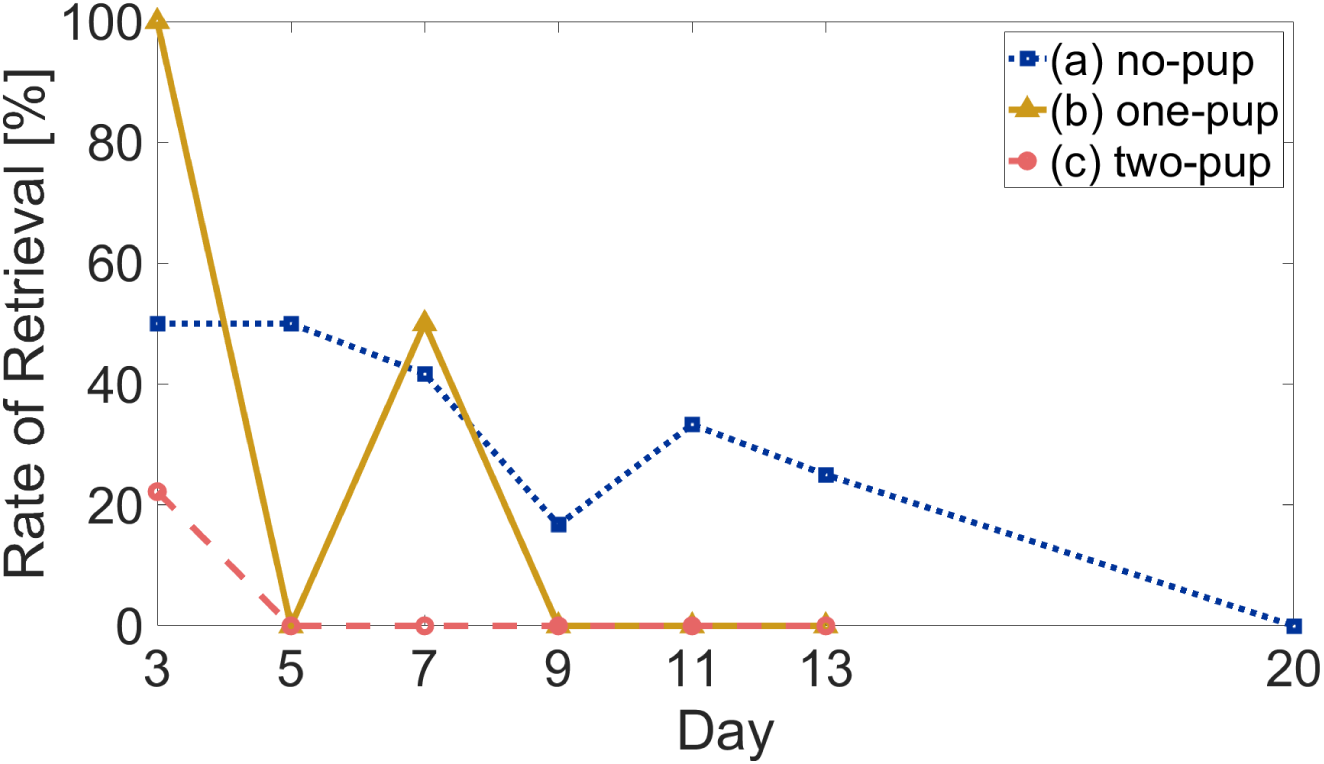
Changes in maternal retrieval rate with pup age in Experiment I. The proportion of trials in which mothers retrieved their own pups is plotted against postnatal age for the three conditions: a) no-pup (blue, n=6), b) one-pup (yellow, n=2), and c) two-pup (pink, n=3). Data on day 20 were obtained only for condition c.

**Table 1.**
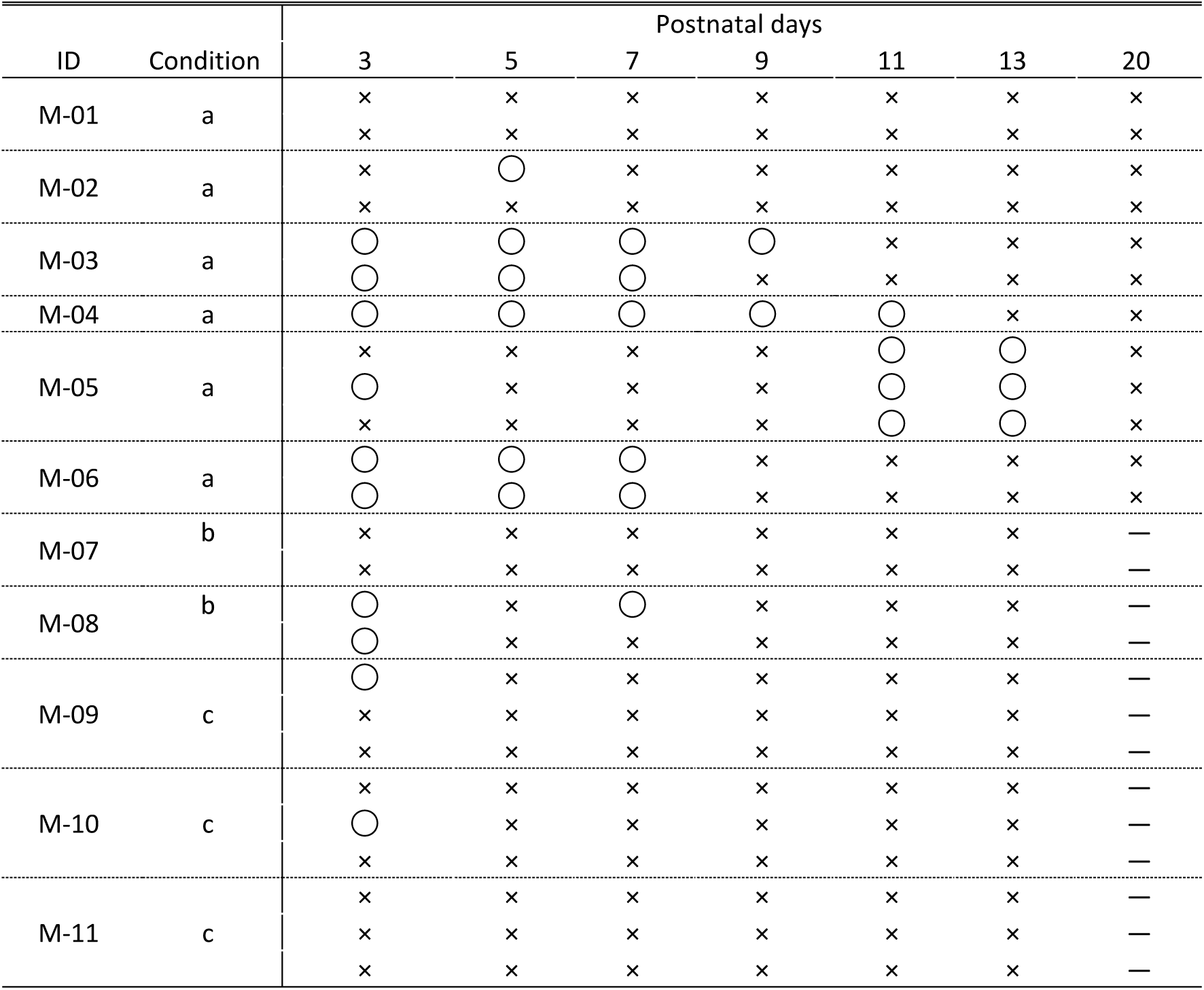
Retrieval results for each mother across pup age in Experiment I. Presence (○) or absence (×) of retrieval behavior is shown by postnatal day and experimental condition (a: no-pup, b: one-pup, c: two-pup). A dash (―) indicates trials that were terminated before completion due to pup movement.

Ambient room temperature was approximately 25°C. Pup skin surface temperatures recorded through the fur using a thermographic camera showed distinct developmental differences. On day 3, when separated from the mother, pup skin surface temperatures fell from below 30°C toward ambient temperature, indicating an inability to maintain body temperature independently. In contrast, on day 11, skin surface temperatures remained stable between 30°C and 34°C despite separation, consistently exceeding ambient temperature, indicating the development of thermoregulatory capacity (Figure S1).

### Experiment II: Playback test

To determine whether the mothers’ retrieval behavior was selectively triggered by the isolation calls of their own pups, we simultaneously presented two playback stimuli: the call of the mother’s own pup from one tweeter, and the call of a non-own pup of the same postnatal age from the other. The mother’s choice of zone was then recorded. Most mothers that entered each zone made physical contact with the tweeter (see Supplemental Video 2).

Figure 5A shows the developmental change in response scores across all trials for five tested mothers. Individual results for each mother are summarized in Table 2. Under the no-pup and one-pup conditions, the scores were high on postnatal day 3 to 9, indicating the mothers responded to the pup calls. Although the scores remained similarly high through day 9, they showed a marked decline between postnatal days 11 and 13. In contrast, under the two-pup condition, response scores remained at 0 across all test days, indicating no response to either playback stimulus. The number of echolocation calls emitted by mothers was significantly higher in trials in which they responded than in those in which they did not (Figure S2).

**Table 2.** Response score results for each mother across pup age in Experiment II. Scores by postnatal day and experimental condition (a: no-pup, b: one-pup, c: two-pup) were assigned as follows: 1 for entering the playback zone, 0 for remaining in the waiting zone, and ― for conditions not tested.

| ID | Condition | Postnatal days |  |  |  |  |  |
| --- | --- | --- | --- | --- | --- | --- | --- |
|  |  | 3 | 5 | 7 | 9 | 11 | 13 |
| M-12 | a | 0 | 0 | 0 | 0 | 0 | 0 |
| M-13 | a | 1 | 1 | 1 | 1 | 1 | 0 |
|  | b | 1 | 1 | 1 | 1 | 0 | 0 |
| M-14 | a | 1 | 1 | 1 | 1 | 1 | 0 |
|  | b | — | 1 | 1 | 1 | 1 | 0 |
|  | c | 0 | 0 | — | — | — | — |
| M-15 | a | — | — | 1 | 1 | 0 | 0 |
|  | c | — | — | 0 | 0 | 0 | 0 |
| M-16 | a | — | — | 1 | 1 | 1 | 1 |
|  | c | — | — | 0 | 0 | 0 | 0 |

Figure 5B presents the latency from playback onset to the mother’s first entry into the playback zone under no-pup condition. The latency increased progressively with pup age.

As shown in Figure 5C, the proportion of approaches to own pup calls was higher than that to non-own pup calls under the no-pup and one pup condition. Under the two-pup condition, mothers remained in the waiting zone for the entire trial (consistent with their baseline tendency to remain stationary in the absence of external stimuli). This indicates that the mother shows a preference for the pup calls.

**Fig. 5.**
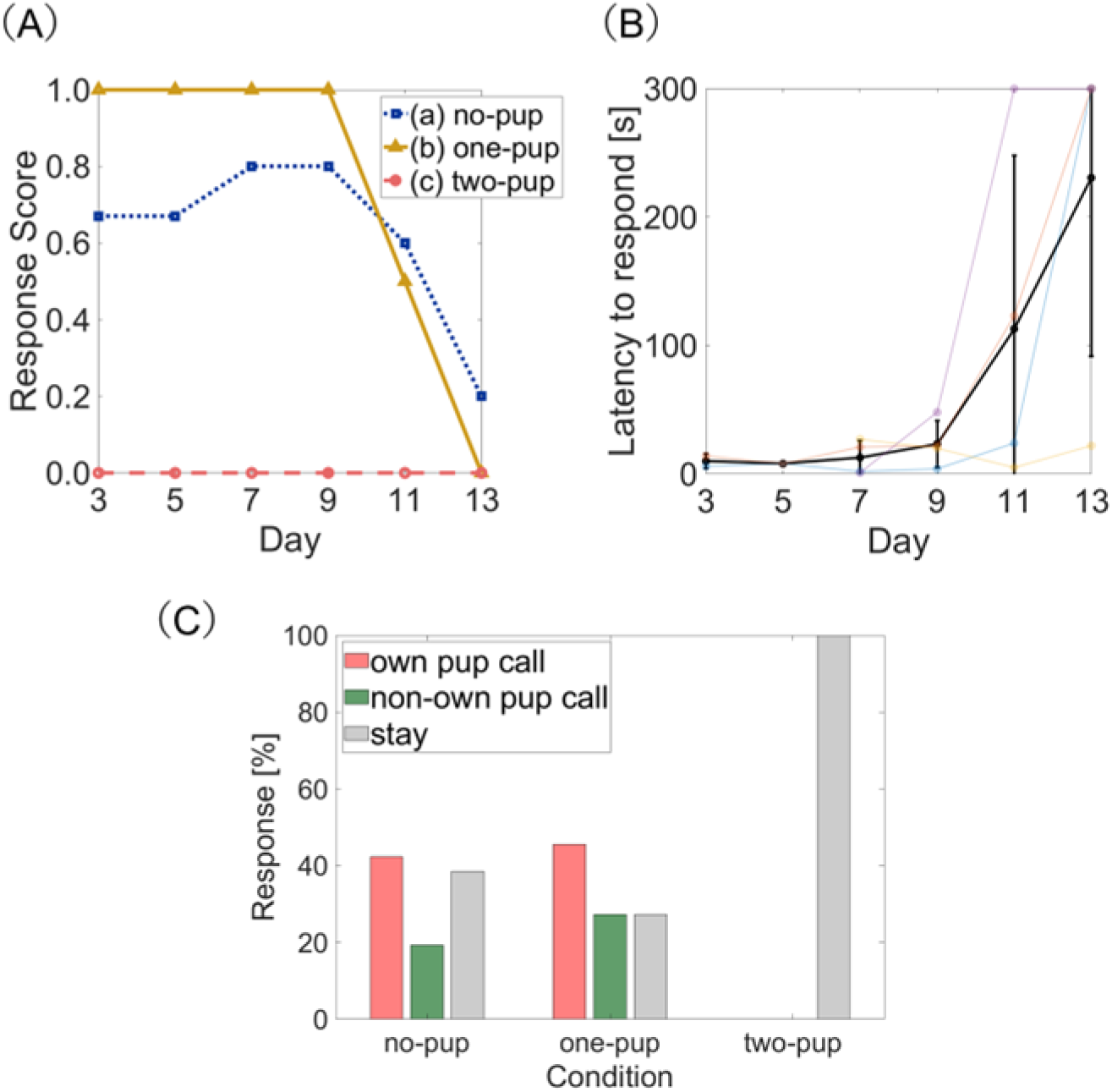
Maternal responses to playback of pup calls across postnatal development. (A) Mean response scores for five mothers under three conditions. Higher scores indicate behavior consistent with higher maternal motivation. A mean of 1 indicates unanimous selection of the mother’s own pup call, whereas a mean of 0 indicates no response to the playback. Sample sizes varied across days and conditions (condition a: n = 3 to 5; condition b: n = 1 to 2; condition c: n = 1 to 2). (B) Latency from playback onset to the mother’s first entry into the zone where the playback (under no-pup condition). Plots show the mean ± SE (black). Colored symbols connected by lines represent individual data each shown in a unique color (n = 5; excluding missing data). (C) Maternal choice responses in the playback experiment. Bars indicate the proportion of trials in which mothers selected the own pup call, non-own pup call, or remained in the waiting zone (stay) under each condition.

Figure 6 shows the mothers’ response scores when playback of pup calls recorded on postnatal day 7 was presented on postnatal days 20 and 35. On the recording day (day 7), mothers under the no-pup and one-pup conditions exhibited high scores, responding to the pup calls. In the no-pup condition, high scores were also maintained on postnatal day 35. In contrast, mothers in the one-pup condition showed no response on postnatal days 20 and 35, resulting in scores of 0. In the two-pup condition, response scores remained at 0 throughout all test days.

**Fig. 6.**
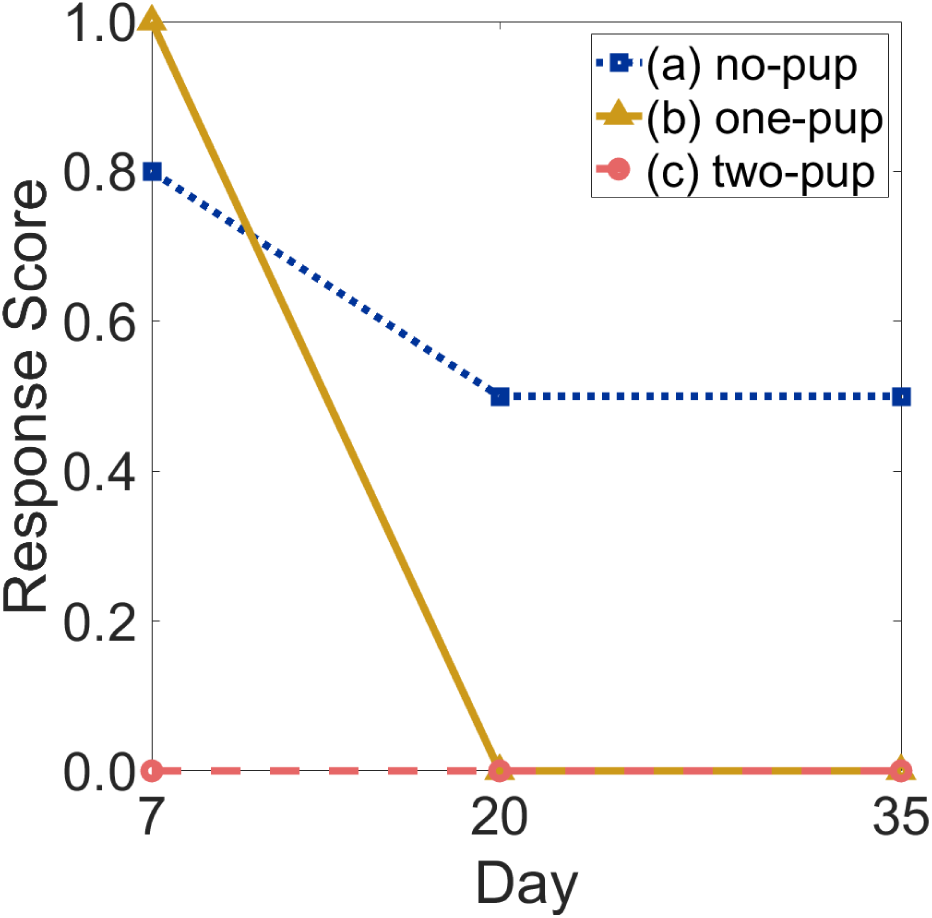
Maternal response to playback of day-7 pup calls (including isolation calls) at later developmental stages. Mean response scores for four mothers are shown for postnatal days 7, 20, and 35 under the three conditions. Mothers without pups maintained high scores even on day 35, whereas those holding pups showed a marked decline. Sample sizes were n = 5 for condition a, and n = 2 for both conditions b and c.

A comparison of the approaching vs. no-approaching model with the GLMMs revealed that the additive model (without interaction between two fixed effects; the number of holdings and the pup age) had the lowest AIC (see Table S5 for all AICs). One of the fixed effects, the number of holdings, exhibited a significant negative effect on retrieval probability (Estimate ± SE = -4.16 ± 1.66, *z* = -2.50, *p* = 0.012) in the selected model. Conversely, the fixed effect of the postnatal days of the mother exhibited no significant effect (*z* = -1.30, *p* = 0.193) in the selected model. Here, the interaction model was excluded from comparative analysis due to its failure to converge.

A comparison of the own vs. non-own selection model using GLMMs showed that the null model (without all fixed effects) had the lowest AIC (see Table S6 for all AICs). And the sole fixed effect model which was consistent with the call rate differences exhibited approximately equivalent AIC (ΔAIC = 0.4). However, no statistical significance was observed for the fixed effect (*z* = 1.20, *p* = 0.231).

### Changes in isolation calls across postnatal age

Figure 7A shows the changes in the number of isolation calls produced by pups across postnatal age. The call rate decreased markedly with increasing pup age: it averaged 8.9 calls s⁻¹ on day 3 but dropped sharply to 0.61 calls s⁻¹ by day 11. As shown in Fig. 7BC, both isolation calls and echolocation precursor calls were produced at similar proportions on days 7 and 9; thereafter, the number of isolation calls decreased while the proportion of echolocation precursor calls increased. By days 11–13, almost all vocalizations were echolocation calls (day 11: 82.3%, day 13: 86.5%).

**Fig. 7.**
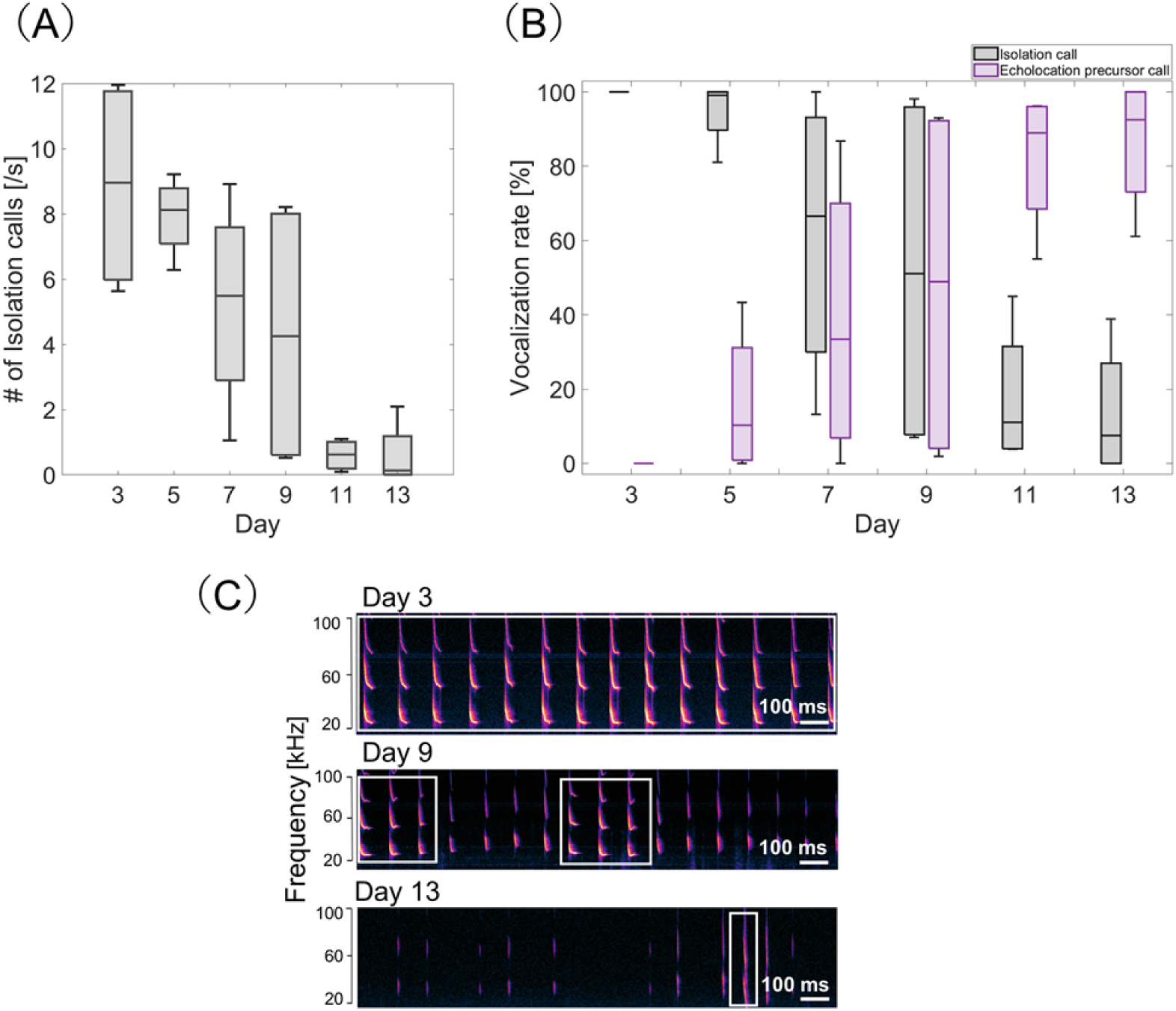
Developmental changes in isolation calls across postnatal day. (A) Number of isolation calls per second. (B) Proportional transition from isolation to echolocation calls with pup development. In both panels, Boxes show the median (center line) and interquartile range; whiskers extend to the most extreme values within 1.5 × IQR, and no outliers were observed. (C) Representative spectrograms of calls recorded on days 3, 9, and 13 illustrate the transition from isolation to echolocation calls; isolation calls are marked by white boxes.

## Discussion

### Dual regulation of maternal retrieval behavior by pup signals and maternal motivational state

In this study, we demonstrated that maternal pup retrieval behavior in *Pipistrellus abramus* is strongly regulated by both pup-derived acoustic signals and the mother’s tactile contact with her pup. GLMM analysis of retrieval probability revealed significant negative effects of both the age of pups and the number of holding pups. Mothers retrieve their pups during the developmental period in which pups produced isolation calls. Furthermore, the decrease of retrieval behavior with the number of holding pups indicates that internal motivational state, shaped by the number of pups in contact with the mother, plays a critical role in the decision to retrieve. The findings indicate that pup retrieval in this species depends on the convergence of two conditions: (i) the presence of isolation calls that signal pup dependence, and (ii) a maternal state in which pup loss intensifies motivational drive. This suggests the presence of a dual-cue mechanism, in which sensory input and internal state collectively dictate maternal behavior.

In Experiment I, maternal response was most frequently observed on postnatal day 3, suggesting that maternal motivation peaks during the immediate postpartum and early neonatal periods. This trend is consistent with previous studies demonstrating hormonal changes accompanying parturition and the existence of a neural sensitive period that facilitates the expression of maternal behavior (Numan and Insel, 2003; Sachser et al., 2020). The reduced responsiveness observed from day 9 in both Experiments I and II may be associated with the loss of this sensitive period. Consistent with this interpretation, analyses of mother’s echolocation call rates (Fig. S2) increased during responsive trials, suggesting greater engagement with the pup stimuli. However, this increase may reflect multiple factors, including enhanced information gathering as well as elevated arousal or agitation. Conversely, the low call rate observed in non-responsive trials likely reflect reduced attention to the presented stimuli.

As pups mature, the disappearance of maternal responses by postnatal day 20 under all conditions likely reflects the pups’ increasing independence associated with the development of thermoregulation (Figure S1), motor coordination, and echolocation abilities (Buchler, 1980). In the playback experiment (Experiment II), the gradual increase in latency for mothers to respond with pup age (Fig. 5B) also indicates a decline in maternal motivation. Thus, as in other mammals, maternal behavior in bats appears to be adaptively expressed only during the early developmental stages when pups are highly dependent on their mothers.

In Experiment II, a GLMM of the approaching behavior toward speakers revealed a significant negative effect was observed for the number of holding pups. This result indicates that the mothers’ behavioral choices varied depending on the situational context, such as the number of pups currently in contact. Physical contact with pups serves as a highly important sensory stimulus for mothers of mammals including human (Lucion and Bortolini, 2014; Stack and Muir, 1992), and ventral somatosensory input elicited by nipple stimulation, etc., in particular, plays an important role in both the initiation and maintenance of maternal responses (Neumann et al., 1993; Stern, 1996). In lactating ewes, the presence of a suckling lamb suppresses HPA-axis responses to psychosocial stress, indicating that mother-infant physical contact can support maternal physiological and emotional stability (Ralph and Tilbrook, 2016).

These findings, well documented in other mammals, suggest that similar somatosensory mechanisms may also operate in this species.

### Energetic and developmental constraints shaping maternal motivation and retrieval behavior

Reproduction in bats entails high energetic demands, particularly during lactation and thermoregulation of dependent young (Barclay et al., 2003; Hayssen, 1993; Racey, 1982). Producing and rearing multiple offspring simultaneously further increases these energetic requirements (Garbino et al., 2021). In mammals, the number of functional nipples has been proposed to represent a physical and functional upper limit on the number of offspring that can be successfully reared (Rawal, 2019). Because *Pipistrellus abramus* typically gives birth to two or three pups but possesses only two nipples, the condition of holding two pups likely approaches the mother’s physiological limit. Indeed, in captivity, mothers are frequently observed carrying two pups simultaneously. When the number of pups being held falls below the number of functional teats—representing a “pup-loss” condition—the mother’s motivational drive is likely elevated, thereby enhancing retrieval behavior. This hypothesis aligns with parental investment theory, which posits that parental behavior is optimized to maximize the survival value of remaining offspring (Clutton-Brock, 1991; Trivers, 1972).

In the playback experiment using vocalizations recorded on day 7 (including isolation calls), mothers in the one-pup condition showed a complete decline in their response scores to zero on postnatal days 20 and 35. This indicates that mothers did not merely respond specifically to their own pup’s isolation calls, but that their behavioral responses were also strongly influenced by the contextual cue of whether the calls were emitted during a developmental stage when pups still required maternal retrieval. In contrast, mothers in the no-pup condition maintained high response scores even on day 35. This is likely because for the mother, the internal motivational state induced by complete pup loss takes precedence in eliciting maternal behavior. Under this circumstance, the internal motivational drive associated with pup loss may have outweighed the developmental appropriateness of the vocal stimuli. Therefore, it is likely that the mothers selectively responded to these calls as rescue signals, even though such vocalizations would normally not be produced by pups at that developmental stage.

### Developmental transition and signaling function of pup isolation calls

In both Experiments I and II, pups frequently produced isolation calls on postnatal day 3, and mothers exhibited the strongest responses during this period (Figs. 4 and 5A). In contrast, by days 11–13, isolation calls were rarely observed and had transitioned into echolocation calls (Figs. 7 and 8). This developmental shift coincided with a marked decline in maternal responses in both experiments, supporting the idea that isolation calls serve as the primary stimulus eliciting maternal behavior, and that their occurrence is strictly dependent on pup developmental stage. Isolation calls were produced only during the early immature phase, suggesting that they function as acoustic indicators of developmental immaturity.

Moreover, mothers showed stronger responses and shorter response latencies when pups produced calls at higher rates, indicating that increased calling frequency enhances the signaling function of the calls and more effectively triggers maternal retrieval behavior. Consistent with this interpretation, retrieval latency was negatively correlated with pup vocalization rate under the no-pup condition (Pearson’s *r* = -0.46, *p* =0.044; Fig. S3). In rats, previous studies have shown that pups emit ultrasonic vocalizations more frequently and for longer durations immediately after maternal contact or during separation (Hofer et al., 2001; Shair et al., 2015). These findings suggest that an increase in calling rate reflects heightened motivation for maternal contact or greater separation-induced distress, a phenomenon that may also occur in *Pipistrellus abramus*. In Experiment II, the number of vocalizations (i.e., pup call rate) did not significantly affect sound-source selection, according to the GLMM results. This suggests that bats do not simply choose pups that emit calls more frequently but instead rely on additional acoustic or sensory cues to selectively retrieve their own pups. Taking together, these findings indicate that isolation calls not only reflect developmental immaturity but also function as behavioral signals of the pup’s negative emotion state. The intensity of their calls (i.e. call rate) may influence maternal responsiveness, whereas selective retrieval likely depends on additional cues.

The isolation calls of *P. abramus pups* can thus be interpreted as a form of distress vocalization within the context of a protest response to separation, sharing common characteristics with human infant crying and the separation calls of other mammals (Hofer, 1996; Newman, 2007). Such distress calls are considered evolutionarily conserved innate signals that elicit parental approach behavior across mammalian species.

## Conclusion

This study provides the first systematic investigation of maternal behavior in the Japanese house bat (*Pipistrellus abramus*) from behavioral and sensory-physiological perspectives. Our results show that both auditory cues from pup isolation calls and somatosensory input through physical contact jointly and flexibly regulate maternal motivation for retrieval. Maternal behavior in this species thus represents a highly adaptive and precisely tuned system integrating multiple sensory modalities to produce context-dependent responses.

In this nocturnal species, which relies heavily on auditory cues under dark conditions, tactile contact becomes the most prioritized information source during the early neonatal period, when mothers constantly hold their pups. Accordingly, the absence of tactile feedback likely signals an immediate crisis of pup loss, rapidly activating maternal responses.

The present findings demonstrate that maternal behavior in *P. abramus* changes adaptively and instantaneously in response to the pup’s condition, offering key insights into innate mechanisms of mammalian maternal care. Because this short-lived species (Funakoshi and Uchida, 1982) invests intensively in offspring, it provides a valuable comparative model—distinct from rodents—for understanding the diversity of maternal regulation and parental investment strategies, contributing to the broader understanding of the evolution and adaptation of mammalian maternal behavior.

## Acknowledgements

We thank Yuna Nishiuchi, Keisuke Kani, and Ryota Yamanaka for their invaluable assistance during the behavioral experiments and data collection; Yuichi Mizutani for helpful advice in statistical analyses.

The authors declare no conflict of interest.

## Data availability

The acoustic recordings, behavioral data, and processed datasets supporting the results of this study are available at figshare (https://doi.org/10.6084/m9.figshare.31007452). Other raw data supporting the findings of this study are available from the corresponding author upon reasonable request.

