## supplementary information for "Tactile pup loss and acoustic signal enhance selective maternal retrieval behavior in echolocating bats, *Pipistrellus abramus*"

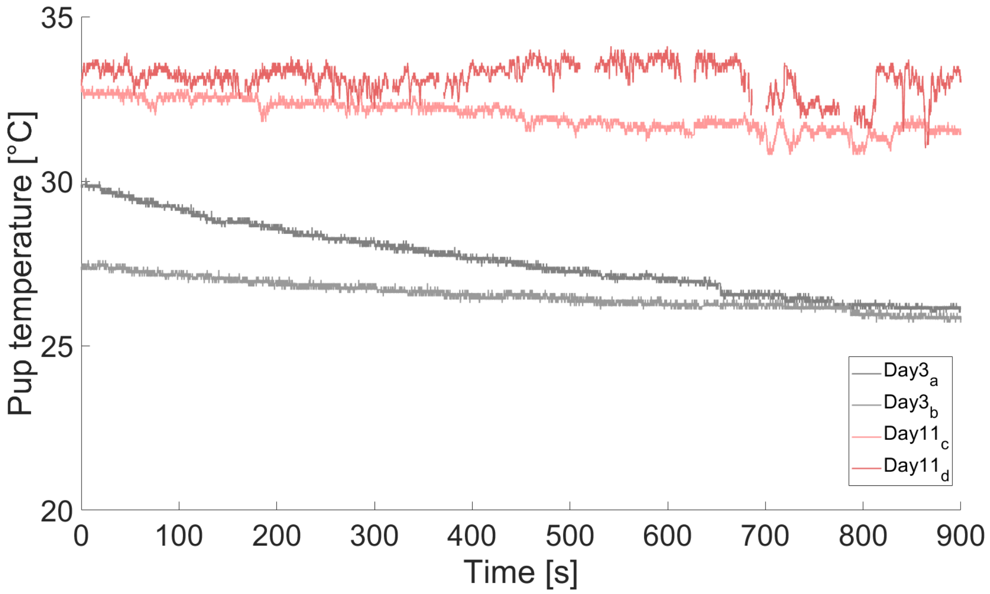


**Fig. S1.** Time-series changes in the surface body temperature of four pups, recorded using a thermographic camera (Xi400, Optris, Germany) positioned 70 cm above the pups on days 3 and 11. Ambient temperature was approximately 25°C. Pup surface temperatures were recorded through the fur using thermography. On day 3, surface temperatures declined toward ambient (approximately 25°C) after separation from the mother, whereas on day 11 they were maintained at 30–34°C remained consistently above ambient.

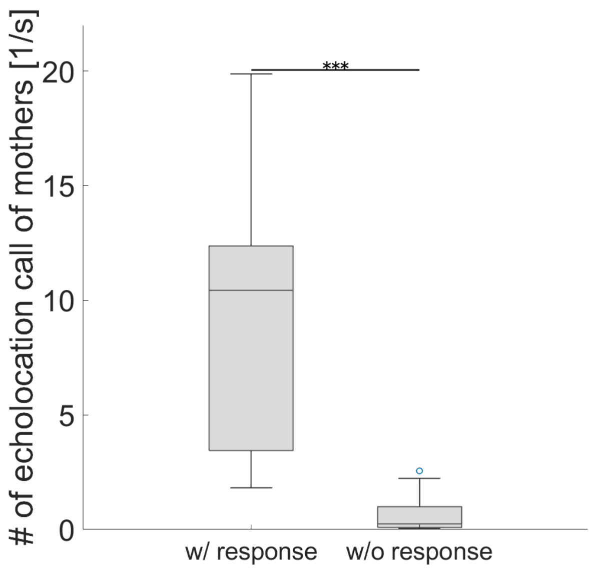


**Fig. S2.** The number of maternal echolocation calls when they showed a response and when they did not in the playback test. They produced more echolocation calls in response to the playback audio, possibly indicating increased efforts to gather information from the surrounding environment. Statistical differences were determined using the Wilcoxon rank-sum test ($W = 364$, ***: $p < 0.001$).

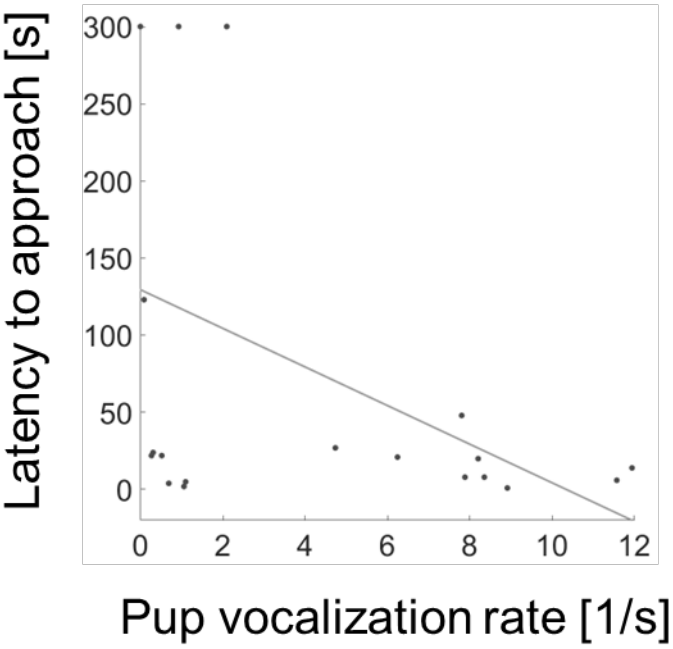


**Fig. S3.** Relationship between response latency and the pup vocalization rate under the no-pup condition in the playback experiment (Experiment II). Each point represents a single trial. The solid line indicates the linear regression. Response latency was negatively correlated with pup vocalization rate (Pearson's $r$ = -0.46, $p$ = 0.044). Latency values were capped at 300 seconds.

**Table S1.** Summary of the pup retrieval test (Experiment I).

Details of mothers tested under each condition (no-pup, one-pup, two-pup), litter size, and the number of trials conducted per day. Applicable conditions are indicated by the number of trials, and non-applicable conditions are indicated by a dash (—).

| ID | Number of births | Number of tests per day | a) no-pup | b) one-pup | c) two-pup |
|---|---|---|---|---|---|
| M-01 | Two | Twice | 14 | — | — |
| M-02 | Two | Twice | 14 | — | — |
| M-03 | Two | Twice | 14 | — | — |
| M-04 | Two | Once | 7 | — | — |
| M-05 | Three | Three | 21 | — | — |
| M-06 | Three | Twice | 14 | — | — |
| M-07 | Two | Twice | — | 12 | — |
| M-08 | Two | Twice | — | 12 | — |
| M-09 | Three | Three | — | — | 18 |
| M-10 | Three | Three | — | — | 18 |
| M-11 | Three | Three | — | — | 18 |
| Total trials | | | 84 | 24 | 54 |

**Table S2.** Comparison of retrieval behavior between biological and non-biological pups in Experiment I (no-pup condition).

Presence (◯) or absence (×) of retrieval behavior is shown. Non-applicable conditions are indicated by a dash (—). Each mother's responses toward her own and non-own pups are listed according to the corresponding postnatal days.

| ID | Postnatal days of mother | Own pup | Non-own pup |
|---|---|---|---|
| M-03 | 13 | × | × |
| M-04 | 11 | ◯ | × |
| | 13 | × | × |
| M-05 | 11 | ◯ | × |
| | 13 | ◯ | × |
| | 13 | ◯ | × |
| M-06 | 9 | × | × |
| | 10 | — | × |
| | 11 | × | — |
| | 13 | × | × |
| Total trials | | 9 | 9 |

**Table S3.** Summary of the playback test (Experiment II).

Experimental conditions for each mother (number of pups held, litter size, and test frequency). Applicable conditions are indicated by the number of trials, and non-applicable conditions are indicated by a dash (—). For mother ID M-14, one pup died during the experimental period; therefore, the number of daily tests was reduced from three to two, and the applicable conditions changed accordingly.

| ID | Number of births | Number of tests per day | a) no-pup | b) one-pup | c) two-pup |
|---|---|---|---|---|---|
| M-12 | One | Once | 6 | — | — |
| M-13 | Two | Twice | 6 | 6 | — |
| M-14 | Three | Three | 1 | 1 | 1 |
| | | Twice | 5 | 4 | 1 |
| M-15 | Three | Twice | 4 | — | 4 |
| M-16 | Three | Twice | 4 | — | 4 |
| Total trials | | | 26 | 11 | 10 |

**Table S4.** Comparison of the AIC of the retrieval behavior GLMM in the Experiment I.

| Model | Fixed effects | Random effects | AIC | ΔAIC |
|---|---|---|---|---|
| MR1 | Holdings + PupAge | (1 + PupAge \| MomID) | **107.6** | **0** |
| MR2 | Holdings × PupAge | (1 + PupAge \| MomID) | 108.9 | 1.3 |
| MR3 | PupAge | (1 + PupAge \| MomID) | 112.6 | 5.0 |
| MR4 | Holdings | (1 \| MomID) | 139.1 | 31.5 |
| MR5 | Null | (1 \| MomID) | 142.5 | 34.9 |

**Table S5.** Comparison of the AIC of the approaching behavior GLMM in the Experiment II.

| Model | Fixed effects | Random effects | AIC | ΔAIC |
|---|---|---|---|---|
| MA1 | Holdings + MomPND | (1 + MomPND \| MomID) | **45.9** | **0** |
| MA2 | Holdings | (1 \| MomID) | 53.7 | 7.8 |
| MA3 | MomPND | (1 + MomPND \| MomID) | 65.0 | 19.1 |
| MA4 | Null | (1 \| MomID) | 65.6 | 19.7 |

**Table S6.** Comparison of the AIC of the own selection GLMM in the Experiment II.

| Model | Fixed effects | Random effects | AIC | ΔAIC |
|---|---|---|---|---|
| MS1 | Null | (1 \| MomID) + (1 \| OwnPupID) + (1 \| OtherPupID) | **38.6** | **0** |
| MS2 | CallDifferences | (1 \| MomID) + (1 \| OwnPupID) + (1 \| OtherPupID) | 39.0 | 0.4 |
| MS3 | Holdings | (1 \| MomID) + (1 \| OwnPupID) + (1 \| OtherPupID) | 40.5 | 1.9 |
| MS4 | MomPND | (1 + MomPND \| MomID)<br>+ (1 \| OwnPupID) + (1 \| OtherPupID) | 41.1 | 2.5 |

**Video S1.** Sample video of retrieval scene in Experiment I.

**Video S2.** Sample video of entering playback area in Experiment II.
